# A hierarchical orthology framework reveals viral carbohydrate-active genes across the global virosphere

**DOI:** 10.64898/2026.08.04.742344

**Authors:** Lingjie Meng, Ruixuan Zhang, Cristina De Castro, Ikuo Uchiyama, Minoru Kanehisa, Hiroyuki Ogata

## Abstract

Carbohydrate-active enzymes (CAZymes) shape virus–host interactions by modifying virion structures, host surfaces and extracellular glycans. However, the diversity and evolutionary origins of viral carbohydrate-active enzymes remain poorly understood, partly due to limited viral protein annotations. To address this, we present VirGenes, a database of viral orthologous groups constructed from the KEGG viral gene dataset. VirGenes uses a hierarchical framework that integrates sequence similarity, remote homology, and structural similarity to support evolutionary and functional analyses of viral proteins. By screening the sequence space of VirGenes, we identified 558 CAZyme-associated gene clusters spanning 102 CAZyme families, revealing particularly enriched repertoires in dsDNA viral lineages. Two bacteriophage families, *Kleczkowskaviridae* and *Pootjesviridae*, encoded more than 10 CAZymes per genome, followed by *Mimiviridae*, a representative family of eukaryotic giant viruses. Phylogenetic analyses systematically revealed divergent evolutionary histories of viral carbohydrate-active genes, including frequent horizontal transfer of endolysin genes from bacteria, which likely represents a viral strategy in the ongoing evolutionary arms race with their cellular hosts. Within the structural space of VirGenes, a large number of viral genes were found to contain CAZyme-like folds despite more than 85% of them lacking detectable sequence similarity to annotated CAZyme sequences. Notably, numerous hypothetical sequences from giant viruses exhibited glycoside hydrolase–like five-bladed β-propeller folds. Overall, by integrating sequence, structural and functional evidence, we show that viral carbohydrate-active systems exemplify how distributed innovations, constrained by ancient folds, collectively build the functional complexity of the global virosphere. VirGenes is publicly accessible at https://www.genome.jp/vogdb/.

## Introduction

Life is a well written symphony of biological molecules, where nucleic acids compose the fundamental scripts, proteins generate the basic functions, and sugars add specificity, diversity and complexity. Sugar-involved processes, including the synthesis, catabolism, and evasion from immune systems, are mediated in part by carbohydrate-active enzymes (CAZymes)^1^. These processes represent the most diverse and significant post-translational modification across all domains of life. For viruses, sugars play essential roles in many processes, such as modulating host-virus interactions and improving the structural integrity of viral particles^2–4^. However, for a long time, viruses were thought to exclusively rely on host sugar-modification machinery to decorate their proteins^5,6^. This paradigm is true until recent discoveries found some viruses encoding their own CAZymes. Specifically, giant viruses were found to have an unexpected repertoire of genes involved in carbohydrate metabolism and glycan synthesis^7–9^, suggesting a more autonomous and complex viral glyco-biology than previously recognized.

Yet, our understanding of viral carbohydrate-active systems remains limited. This knowledge gap is due to the fragmented experimental data and the lack of a systematic bioinformatic framework. Most glyco-related viral genes have been identified in just a few model cultivation systems^10–12^, whereas progress that connects experimentally verified viral carbohydrate-active enzymes to global viral diversity should employ systematic computational analyses. However, previous viral orthologous detection methods relied heavily on sequence similarity, making them ineffective against the rapid evolution and high variability of viral proteins. Because viral proteins can be highly divergent and represented in limited experimental systems, sequence-based searches alone may miss remote homologs that retain conserved structural folds^13,14^. Recent advances in protein structure prediction now allow the identification of viral homologs and functions across fold space. We therefore considered the detection of viral carbohydrate-active enzymes could be improved by building a comprehensive viral orthologous group database that spans both sequence and structure similarity space.

Here, we present VirGenes, a unified database of viral orthologous groups built from the KEGG viral gene dataset^15,16^, with a focus on detecting viral carbohydrate-active genes. VirGenes classifies viral protein families across multiple levels of homology, including <u>V</u>iral <u>S</u>equence <u>C</u>lusters (VSCs), <u>V</u>iral <u>R</u>emote <u>C</u>lusters (VRCs) and <u>V</u>iral <u>F</u>old <u>C</u>lusters (VFCs).

Here, “orthologous groups” is used in an operational sense to describe functionally coherent groups of genes, rather than strict orthologs inferred through speciation-aware analysis. The clustering parameters were optimized to realize high consistency with KEGG Orthology (KO) assignments, followed by additional cluster refinement. The sequence-based clusters (VSC) were further grouped by remote homology detection (VRC) as well as structural similarities (VFC). This hierarchical design allows both closely related and highly divergent viral homologs to be connected within a common framework. Using VirGenes, we systematically characterize the distribution, evolutionary origins and structural diversity of viral CAZyme repertoires across the global virosphere. Here, we use “carbohydrate-active genes” to refer to genes encoding CAZyme proteins, including glycosyltransferases (GTs), glycoside hydrolases (GHs), polysaccharide lyases (PLs), carbohydrate esterases (CEs), auxiliary activity enzymes (AAs) and carbohydrate-binding modules (CBMs), capturing enzymes directly involved in glycan synthesis, modification, degradation and recognition.

## Results

### Construction, refinement and assessment of a hierarchically structured VirGenes

In this study, we first developed a hierarchical framework for grouping viral genes (VirGenes), which is roughly summarized in a schematic workflow (Fig. S1). Briefly, the workflow began with the collection of viral genes and genomes from the KEGG database (www.kegg.jp), accompanied by other resources, such as the Virus-Host DB (www.genome.jp/virushostdb/) and the ICTV’s VMR (ictv.global). The original viral gene set included ORFs that encode long polyprotein precursors, which can induce heterogeneous clusters. To overcome this problem, we added a preprocessing step before clustering by identifying and segmenting putative polyprotein precursors into mature peptide-like regions. According to the keyword search and alignment against KEGG’s curated mature peptide set, a set of 1,048 sequences were labeled as “polyproteins” (Fig. S2a, b). Using these labels, we evaluated our polyprotein-processing approach (Fig. S2c). Approximately 90% of the labeled polyproteins were successfully segmented into multiple regions. A case study showed that the well-characterized coronavirus polyprotein ORF1ab was partitioned into more than 14 fragments, closely matching the number of proteins established experimentally (Fig. S2d). Notably, the RNA-dependent RNA polymerase was isolated from the polyproteins, enabling downstream clustering of this individual gene. In total, 4,958 ORFs, recognized as hypothetical polyproteins, in the original dataset were split into 26,738 distinct viral genes. Overall, 703,080 viral genes were processed in the clustering step.

The first layer of clustering was rooted in the sequence space. Sequence clustering was performed using sequence similarity networks and the MCL algorithm, yielding 63,216 viral sequence clusters (VSCs), including 42,297 clusters with at least three members. The intra-group median sequence identity varied across VSCs, with a minimum of 23.68% and an overall median of 61.32% (Fig. S3). Then these raw clusters were further refined with several steps. First, intra-cluster centralities were calculated, and centrality thresholds were applied to remove outlier sequences from each VSC. Additional outliers were then removed based on sequence length. Second, an iterative HMM-profile reassignment step was used to ensure that each retained gene was most strongly associated with the conserved domain profile of its assigned VSC. The refined VSCs were subsequently clustered according to similarities between their conserved domain profiles, generating 3,898 Viral Remote Clusters (VRCs). Functional annotations were assigned to VSCs and VRCs using a majority-rule principle and were further refined through manual curation.

Both VSCs and VRCs represent the sequence space in the VirGenes. To map the sequence space to fold space, protein folds were predicted or integrated from available structural data for each VSC. Then an all-against-all fold alignment was performed, yielding 5,290 fold clusters (VFCs) with more than one VSC. Gene-level KO annotations were integrated into consensus annotations at the VSC level, whereas KO association profiles were derived for VRCs and VFCs from their constituent VSC annotations. Cross-references between VirGenes and KEGG VOG^15^ were established through shared member genes.

For biological context, we looked into the taxonomic distribution of VirGenes. At the VSC level, a clear gradual decrease in VSC content similarity was observed across all viral realms as taxonomic rank increased (Fig. 1a), and members of the same genus showed highly similar gene content (Jaccard index > 0.7). The rarefaction plot showed that despite the number of VSCs found in viral realms was largely different, the discovery of VSCs did not reach a plateau for any of the viral realms (Fig. 1b). The distribution of VSCs across taxonomic groups was highly uneven so that most clusters were restricted to a single taxonomic group. At the same time, however, a subset of VSCs occurred across multiple genera, families, and orders, reflecting both broader gene sharing and the presence of conserved core gene sets within lower-ranked groups (Fig. 1c). The most broadly distributed VSCs were enriched in nucleotide metabolism and genome replication functions, such as dUTPase, thymidylate synthase, RNaseH and ribonucleotide reductase. Despite the wide taxonomic distribution of specific genes, VSC members from different taxa displayed sharp genetic divergence, even at the family level (Fig. 1d).

**Figure 1.**
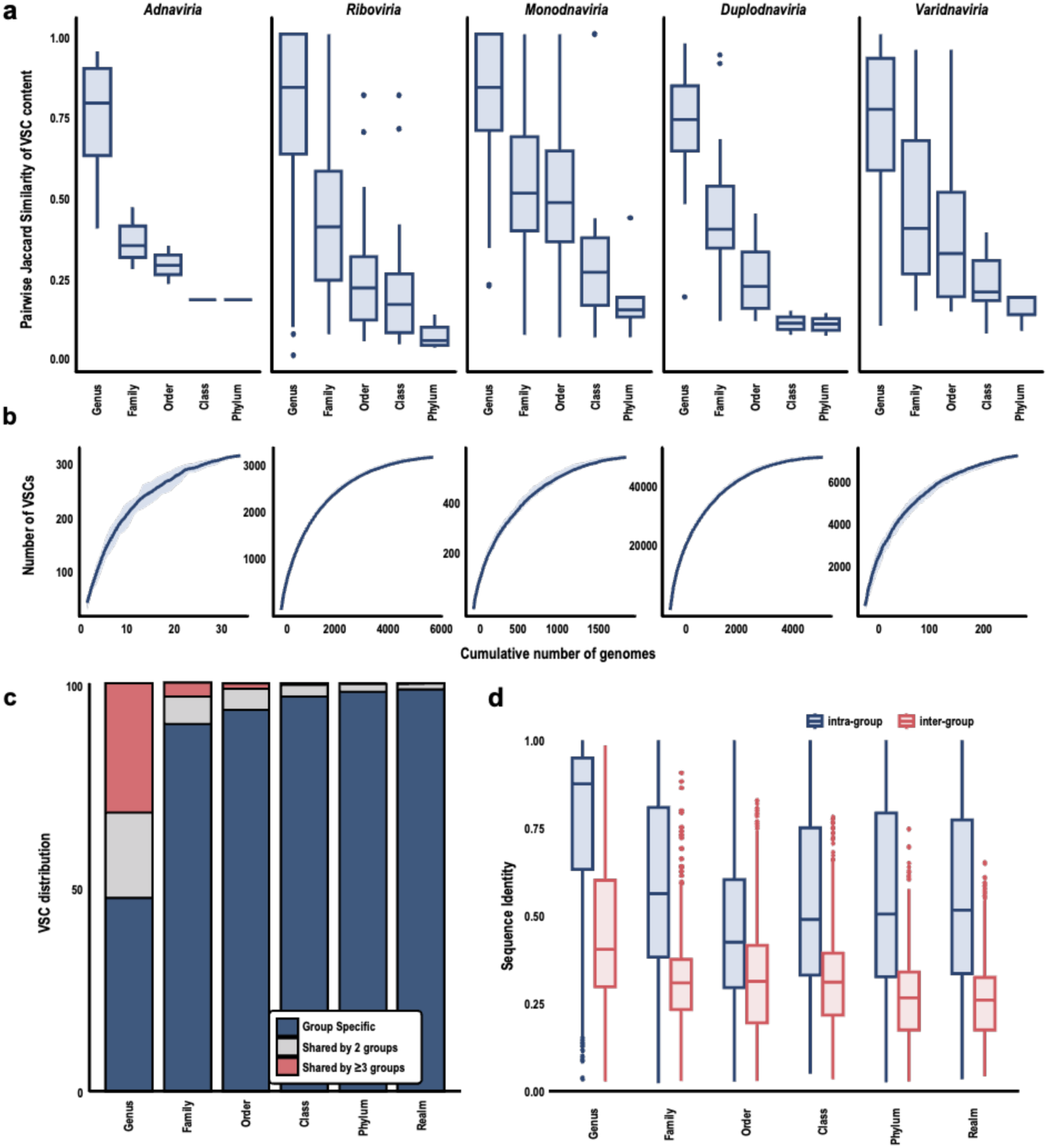
Global landscape of the VIRGENES hierarchical framework. a,VSC compositional patterns across the five viral realms at increasing taxonomic ranks. **b**, Rarefaction analysis of VSC discovery as a function of the number of genomes sampled. **c**, Distribution of VSCs across viral taxonomy groups. **d**, Pairwise sequence similarity across taxonomic levels.

#### Widespread viral CAZyme repertoires in the virosphere

We leveraged the constructed database and mapped viral sequences to CAZyme sequences. We first performed sequence alignment to CAZyme sequences, then ran an HMM search to improve detection sensitivity (Fig. 2a). VSCs showed high intra-group homogeneity in CAZyme gene content. In most cases, a cluster either had no annotated members or most of its members mapped to the same CAZyme family (Fig. S4). Based on this observation, when more than half of the genes of a VSC had hits against the same CAZy family, we assigned that CAZyme label to the entire VSC. This workflow allowed us to define the CAZyme genes with the conserved sequence information present in VirGenes (CAZyme-VSC). In total, we identified 8,807 CAZyme-encoding viral genes from 558 VSCs across 102 CAZyDB families, which exceeds the number of sequences detected by the default HMMER pipeline in dbCAN^17^ or strict alignment-based scans (Fig. 2a). Among the 558 VSCs, 353 had 100% members annotated as CAZymes, indicating a strong consensus in CAZyme identity within VSCs.

**Figure 2.**
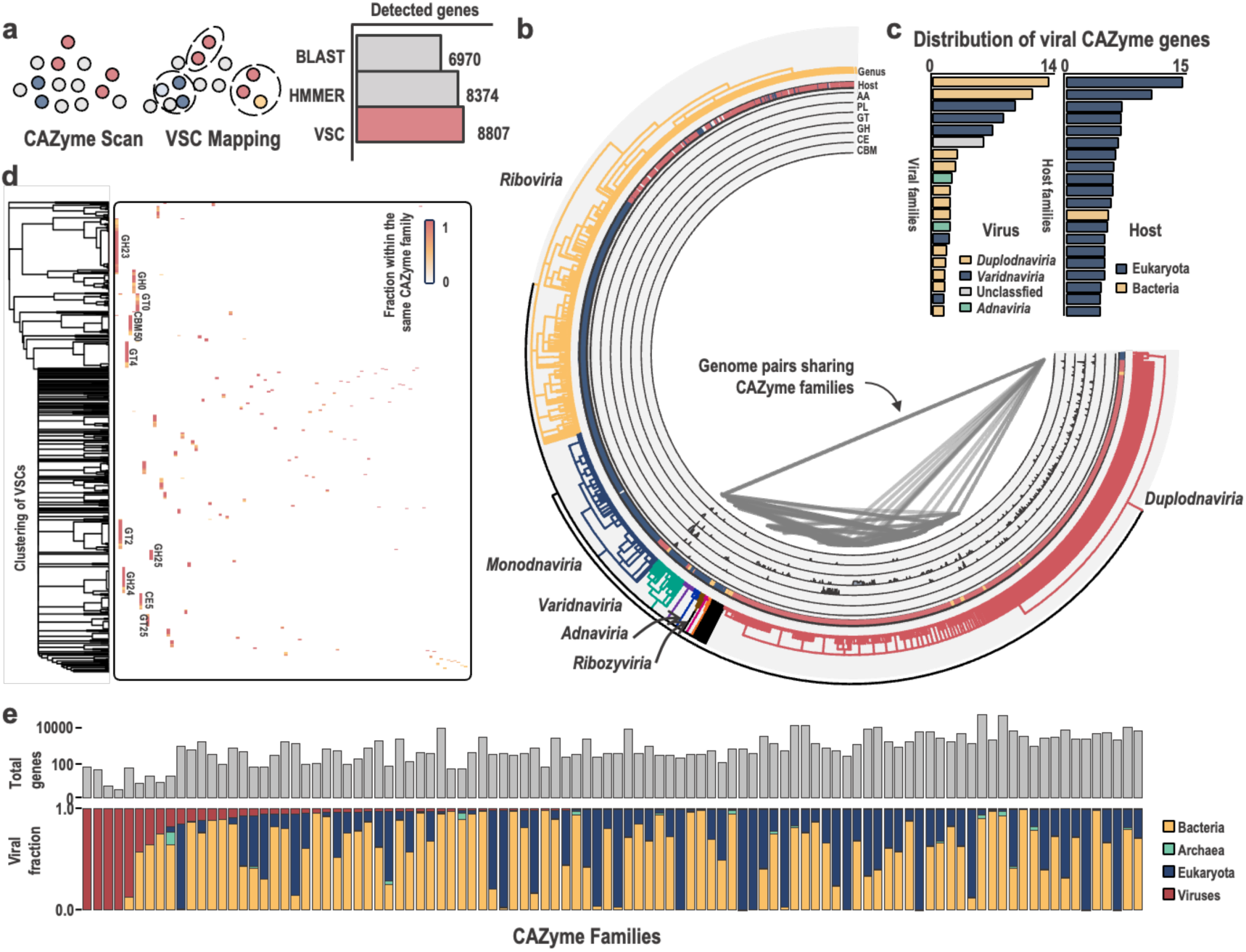
**Viral CAZyme identification and distribution**. **a,** Workflow for detecting viral CAZymes using BLAST, HMMER, and VSC-based annotation. **b,** Taxonomic distribution of CAZyme-containing VSCs across major viral groups. **c,** Distribution of CAZyme families across viral taxa and host types. **d,** Clustering of VSCs by CAZy family. **e,** Composition of CAZy families.

Most CAZyme detections in VirGenes belonged to the realm *Duplodnaviria*. Two phage families of this realm, *Kleczkowskaviridae* and *Pootjesviridae*, had the highest average number of CAZyme genes per genome. However, the total CAZyme reservoir in the realm *Varidnaviria* may be underestimated because of the limited availability of genomes in public databases. *Mimiviridae*, *Phycodnaviridae*, and *Schizomimiviridae* showed high densities of CAZyme-encoding genes, with 380 identified genes across 44 genomes (Fig. 2b, c). By linking these data to the host metadata in VirGenes, we found that several groups of eukaryote-infecting viruses contained particularly rich CAZyme repertoires (Fig. 2b, c). For example, eight viruses infecting hosts in the family Chlorellaceae encoded a total of 120 carbohydrate-active genes. Nucleocytoviruses shared many CAZyme families with both herpesviruses and phages, suggesting that dsDNA viral groups have partially overlapping CAZyme repertoires. Although viral genes in different VSCs generally had distinct cluster boundaries, multiple VSCs were sometimes annotated to the same CAZyme families (Fig. 2d). By incorporating the VRC relationships, we were able to integrate VSCs from the same CAZyme family through conserved domain-level similarities (Fig. S5). We combined the detected viral CAZyme proteins with cellular CAZyme proteins and found that, as expected, most CAZyme families were largely represented by cellular proteins (Fig. 2e). However, several families, including GH83 (sialidase), PL23 (chondroitin lyase), and GH90 (rhamnosidase), were dominated by viral entries, with more than 80% of their members corresponding to viral proteins. GH23 and GH24, both encoding muramidases often associated with endolysin-like peptidoglycan degradation, and GH19 (chitinase) contained the largest numbers of detected viral genes.

#### Evolutionary origins of viral carbohydrate-active genes in the global virosphere

To investigate the origins and evolutionary trajectories of viral CAZyme-encoding genes, we performed phylogenetic analyses across all major CAZyme families including viral sequences. These trees revealed widespread horizontal gene transfer (HGT) events between viruses and cellular lineages (Fig. 3). Among these, gene transfers from bacteria to viruses largely outnumbered those from eukaryotes to viruses. The most extreme case was observed in GH23, where we detected 89 bacteria-to-virus transfer events, compared to only 11 transfers from bacteria to eukaryotes and 9 from bacteria to archaea. Notably, we also identified 12 events of reverse gene flow from viruses back to bacteria, highlighting an underappreciated potential role of viruses as gene reservoirs in microbial ecosystems. A similar pattern was observed in GH24, GH25 and GH19. Considering that the composition of viral genes within these families (e.g., GH23 and GH24) were almost phages, the phylogenies support the idea that the gene gains of glycoside hydrolases are likely being a strategy of viruses for an evolutionary arms race with their cellular hosts.

**Figure 3.**
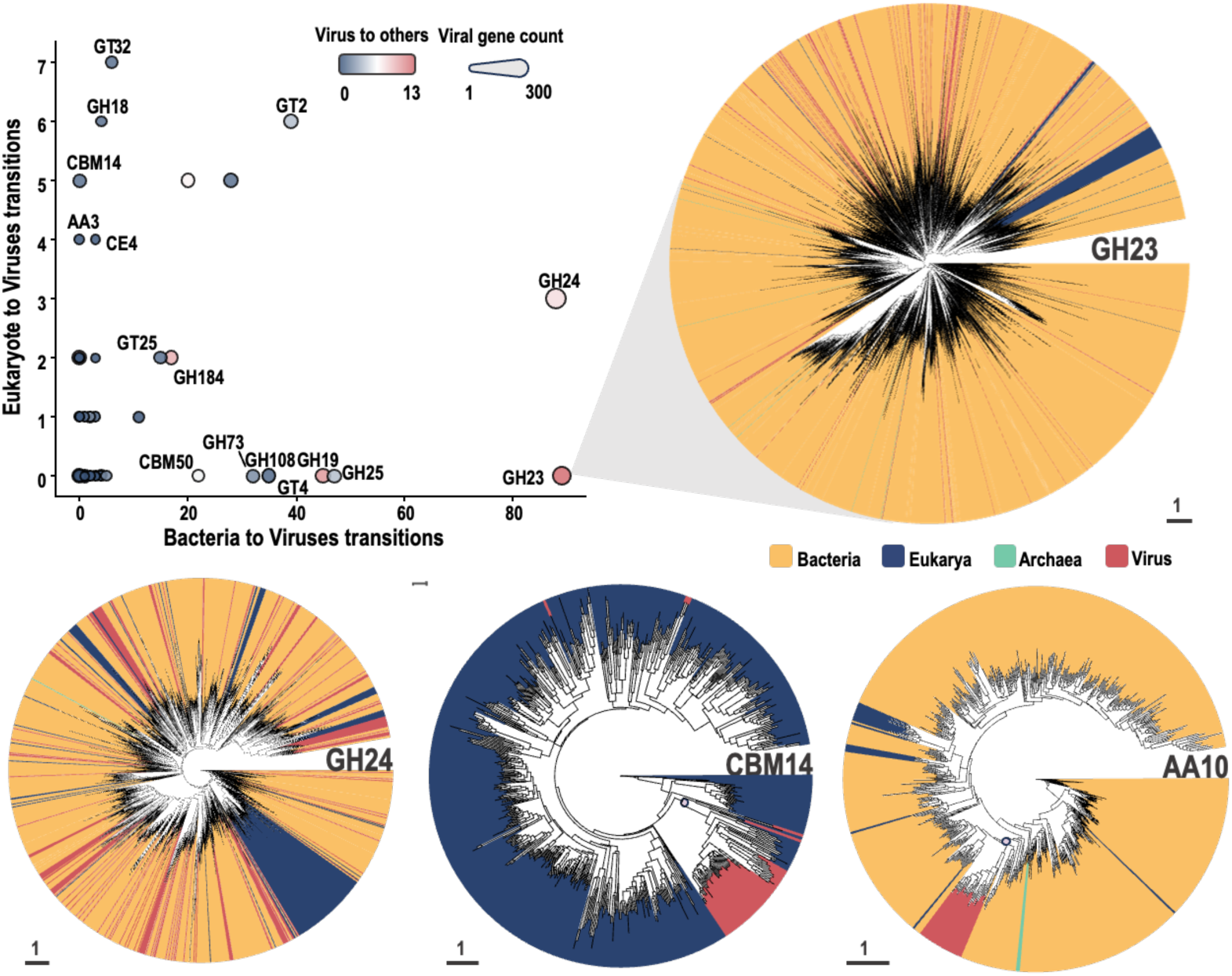
Evolutionary path of viral carbohydrate-active genes. Scatterplot represents transition counts across all gene families, where the x-axis represents bacterial-to-viral transitions, the y-axis represents eukaryotic-to-viral transitions, circle size corresponds to the number of viral members within each family, and color intensity indicates the number of virus-to-host transitions. Phylogenetic trees of representative CAZyme families containing viral genes from the VirGenes, reconstructed maximum likelihood methods. The purple circle indicates the common ancestor of the viral sequences and their sister cellular clade, supported by an ultrafast bootstrap value greater than 95%.

Not all CAZyme families show frequent gene transfer from cellular organisms. For example, in AA10, a family of lytic polysaccharide monooxygenases, phylogenetic evidence suggests a single ancestral acquisition followed by diversification within phages, with little subsequent gene exchange with cellular lineages. We also observed sporadic but notable horizontal gene transfer events from eukaryotes to viruses in certain families. For instance, in CBM14, a clade of baculovirus genes appears to have originated from eukaryotes and subsequently diversified. These viral CAZymes often branch within or adjacent to known eukaryotic clades, and in some cases form well-supported sister groups to eukaryotic sequences, suggesting both recent and ancient horizontal gene transfer events.

#### Identification of glycoside hydrolase–like β-propeller folds

Viral proteins may share structural similarity with functional proteins from cellular organisms while showing little or no sequence homology, due to high mutation rates and divergent evolutionary origins. To systematically uncover potential carbohydrate-active proteins across the virosphere, we performed a global structural alignment between all VSC structural profiles and protein models representing known CAZyme families. In total, 3,548 structural pairings were identified between VSCs (1,302) and CAZyme families (N = 135), forming a network of protein folds. This fold network could be broadly divided into four communities, corresponding to GT folds, parallel β-folds, (α/β)8 folds, and β-propeller folds (Fig. 4a). GH and PL superfamilies formed three major hybrid structural hubs, suggesting a shared architectural scaffold despite their functional and sequence divergence. Of the 1,302 VSCs, only 190 (14.5%) could be linked to CAZyme families based on sequence similarity. The largest number of candidate VSC genes were found in *Duplodnaviria* genomes (N = 3,961), followed by *Varidnaviria* (N = 1,038) (Fig. 4b). However, when normalized by total gene counts, *Varidnaviria* (1.8 genes per genome) emerged as the primary hotspot for novel fold discovery, compared with *Duplodnaviria* (0.4 genes per genome).

**Figure 4.**
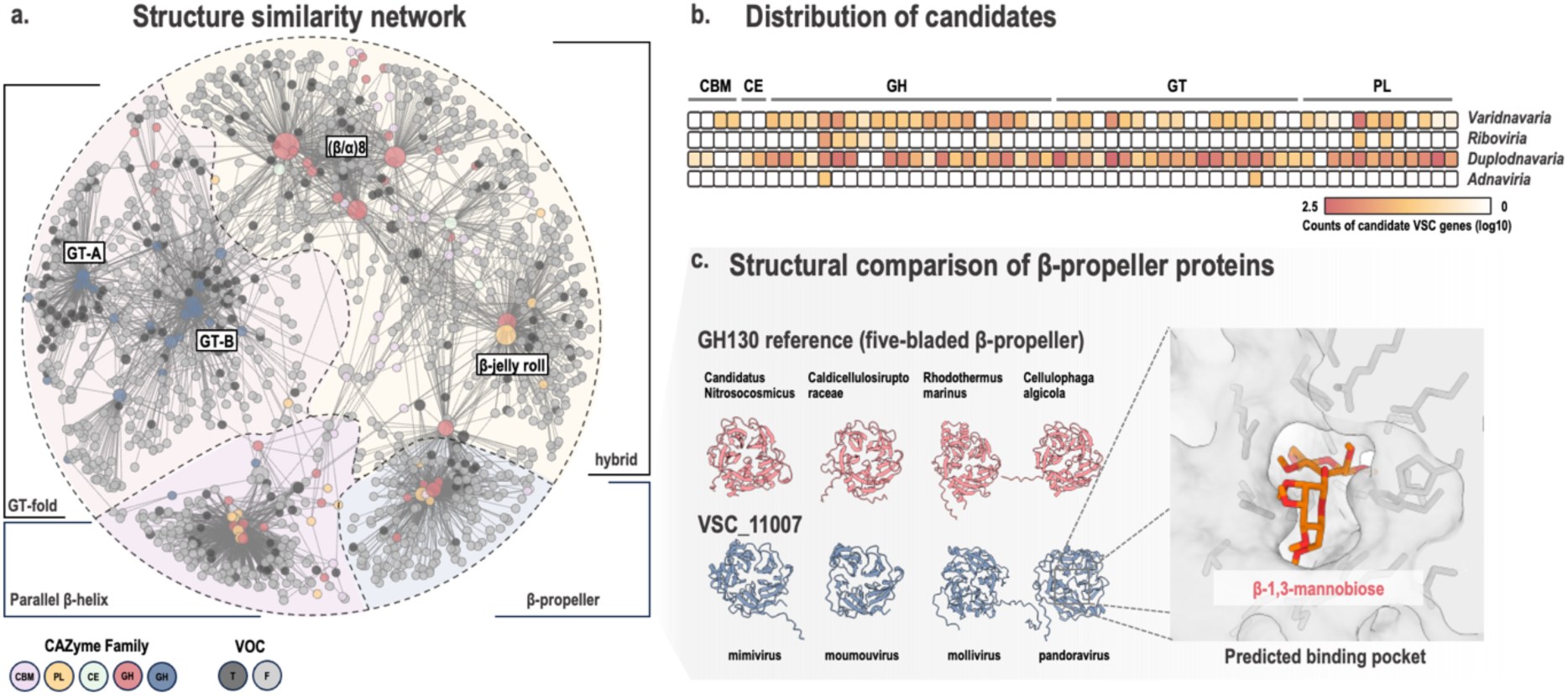
**Detection of the novel viral protein folds**. **a,** Structural similarity network of VSCs and CAZyme families based on Foldseek alignments. **b**, Distribution of candidate CAZyme-like genes across viral realms. Color intensity represents log10-transformed candidate gene counts. **c**, Structural comparison of representative viral β-propeller proteins with GH130 family members. The inset shows the best-scoring docking pose of β-1,3-mannobiose in the highest-ranking predicted binding pocket of a VSC_11007 representative protein.

Among all fold types, the β-propeller community contained many VSCs lacking sequence-based CAZyme annotations (n=34) with high structural similarity (TM-score >0.4) to the GH130 glycoside hydrolase family (five-bladed β-propeller fold) or the PL22 polysaccharide lyase family (seven-bladed β-propeller fold). These VSCs exhibited typical β-propeller topologies but showed no detectable sequence similarity to known CAZyme members. For example, VSC_11007 showed strong structural similarity to multiple GH130 subfamilies despite lacking any detectable sequence similarity. Both the viral proteins in VSC_11007 and GH130 cellular representatives adopted a canonical five-bladed β-propeller fold (Fig. 4c). Additional BLASTP searches against public databases confirmed that members of this VSC are annotated as hypothetical proteins. Notably, this VSC was exclusively composed of amoeba-infecting giant viruses from different viral orders, such as mimiviruses and pandoraviruses. Structural comparisons using the Dali server^18^ further supported the hydrolase-like nature of these viral β-propeller folds. Similarly, other CAZyme families with related folds showed strong structural matches to unannotated VSCs.

To further assess the functional relevance of these novel viral β-propeller proteins, we performed binding-pocket prediction and molecular docking. In both the reference GH130 protein and the VSC_11007 viral protein, the highest-ranking predicted pocket was located within the central groove of the five-bladed β-propeller fold (Fig. 4c). Both pockets contained acidic, basic, and aromatic residues in a similar spatial configuration that may contribute to carbohydrate recognition. Among the tested disaccharides, β-1,3-mannobiose yielded the most favorable mean docking scores for both the reference protein (−6.066 ± 0.005 kcal mol−1) and the viral protein (−6.240 ± 0.005 kcal mol−1), compared with β-1,4-mannobiose, cellobiose, and maltose. This concordant ligand ranking is consistent with a shared preference for β-1,3-linked mannosides and supports the presence of a GH130-like substrate-recognition environment in the viral protein, although experimental validation is required to establish its enzymatic activity and substrate specificity.

## Discussion

Annotating viral proteins remains a major challenge in virology, because viral genes evolve rapidly and often show no detectable homology to cellular genes^19^. Typically, less than ∼30% of viral protein families can be assigned a functional annotation^13,20^. This significantly hinders our understanding of their functions. This limitation represents a fundamental bottleneck in viral biology. Without functional annotation, viral genes cannot be integrated into molecular interaction networks, metabolic pathways, or host-virus interaction frameworks. Moreover, the large proportion of functionally uncharacterized viral genes limits evolutionary inference, as orthology relationships alone provide insufficient insight into selective pressures, functional innovation, and ecological adaptation. Despite the high level of sequence divergence, many viral proteins retain conserved structural architectures^13,21^, underscoring the possibility for annotation frameworks that integrate information across multiple evolutionary scales rather than relying solely on sequence similarity. Building on this principle, in this study, we provide VirGenes with a unified framework that bridges sequence and structure by progressively clustering viral proteins using pairwise similarity, profile-profile comparisons, and structural alignments to connect increasingly remote homologs. Recent resources such as the Big Fantastic Virus Database (BFVD)^22^ and the Viral AlphaFold Database (VAD)^23^ also reveal extensive conservation of viral protein folds. Beyond structure-based clustering, VirGenes integrates KO, which allows mapping viral genes to KEGG pathway resources, and incorporates rich contextual metadata, including host associations and viral taxonomy from resources such as the Virus-Host Database^24^ and ICTV^25^. VirGenes is distinct from the previously reported Virus Orthologous Groups Database (VOGDB)^26^. Although both resources organize viral proteins using sequence and structural relationships, they differ in their source datasets and group-construction strategies. VirGenes is constructed from the KEGG viral gene dataset, incorporating KO-guided optimization and refinement of sequence clusters, whereas the existing VOGDB is based primarily on viral genomes from RefSeq.

VirGenes is also designed to maintain stable accession numbers, ensuring long-term compatibility and enabling seamless expansion as the database grows. Beyond protein grouping, VirGenes is designed as a biology-oriented platform that connects viral protein families with functional evidence, representative structures, viral taxonomy, host distribution, and phylogenetic context. The current web resource allows users to explore individual protein groups from functional, evolutionary, and ecological perspectives.

Future development will further integrate genomic-context information, experimentally supported functions, and other biologically relevant annotations. Overall, VirGenes establishes a foundation for a biology-oriented viral gene resource that connects protein classification with functional, evolutionary, and ecological information.

Using VirGenes, we systematically reveal that carbohydrate-active genes are widespread across the virosphere. Although CAZymes are important to viral infection, such a viral gene-toolkit was overlooked for a long while because most small viruses rely on host carbohydrate-active enzyme systems. Some bacteriophages encode their own endolysins as heat-labile lytic factors during early infection. Many of these endolysins belong to glycosyl hydrolase families such as GH24 and GH25^27,28^. In the 1990s, Paramecium bursaria chlorella virus 1 (PBCV-1) challenged the prevailing view that viruses rely entirely on host metabolism by providing a landmark example of a virus encoding glycosylation-related enzymes, including hyaluronan synthase and enzymes involved in nucleotide-sugar precursor production for hyaluronan biosynthesis^11^. In our data, we identified 8,807 genes from 558 VSCs aligned to 102 CAZyDB families. Notably, even when restricting the analysis to reference viral genomes and using a very conserved threshold, VirGenes still enlarges the detectable repertoire of viral CAZymes. As expected, carbohydrate-active genes are widespread in dsDNA viruses. More specifically, glycoside hydrolase genes were widely detected in phages. These enzymes likely represent one of the most important viral adaptive traits that enable phages to degrade host cell surface polysaccharides, thereby facilitating host recognition, adsorption, and genome entry during infection. Meanwhile, eukaryote-infecting viruses encoded enriched and taxonomically diverse sets of CAZymes involved in host-like glycan synthesis. These patterns indicate that viral CAZyme repertoires are not sporadically distributed but likely represent an adaptive strategy to infect their hosts^4^. The importance of glycosylation is supported by recent experimental work demonstrating that bacteriophage T4 encodes two primary DNA glucosyltransferases with distinct stereochemical specificities, which together enable phage survival under combined host defense pressures, including restriction-modification systems and stereospecific DNA glycosylases^29^. Taken together, our results support the view that viral CAZyme-encoding genes form a host-adaptive gene toolkit shaped by selective pressures.

We then asked how viruses acquire such adaptive strategies. Several GH families, i.e., GH23 and GH24, show clear signatures of repeated horizontal gene transfer (HGT) from bacterial hosts (Fig. 3). Such extensive HGTs have been reported across various GH families in global biological systems, including plant-derived GHs transferred to the whitefly *Bemisia tabaci*^30^. As mentioned above, GH23 and GH24 enzymes could function as endolysins that degrade the bacterial peptidoglycan layer. Their evolutionary dynamics match the expectations of an ongoing phage-bacteria arms race, in which phages continuously acquire and modify GH genes to overcome changes in bacterial cell wall structures. Some studies have shown that endolysins can be horizontally exchanged between phages and rapidly adapt to host through recombination and mutations^31^. In contrast, other CAZyme superfamilies follow distinct evolutionary trajectories, indicating that the selective pressures shaping the evolution of individual CAZymes are different.

A key advantage of VirGenes is its ability to connect sequence space with fold space, which uncovers layers of functional innovation that are entirely inaccessible to sequence-based annotation. Our global fold comparison reveals that most viral genes with folds similar to CAZymes do not have detectable sequence homology. A clear case is that some viruses encode a wide range of β-propeller proteins whose architectures closely mirror those of GH130 and PL22 CAZyme families, although sequence-based annotation still classifies them as hypothetical proteins. These viral β-propellers preserve the canonical blade organization and catalytic pocket geometry, **implying that key mechanistic features have been retained despite deep evolutionary divergence**. Such findings indicate that viruses may harbor previously unrecognized CAZyme-related activities built upon ancient, conserved structural templates. More broadly, this demonstrates the power of a fold-aware framework to illuminate functional signals within viral dark matter and substantially broaden the architectural landscape of viral enzymes.

Together, these findings indicate that viral functional diversity is organized around a set of stable molecular modules with gene clusters, domain architectures, and recurrent structural folds that persist across vast evolutionary distances. The smooth shifts in VSC composition across viral orders and realms further reinforce an emerging view of module-based viral classification, one that complements and extends traditional sequence-derived taxonomies. Although VirGenes remains constrained by uneven genome sampling and the need for experimental confirmation of predicted structures, it establishes a scalable foundation for interpreting viral dark matter. By integrating structural signatures with evolutionary context, VirGenes expands the detectable functional space of viral proteins and provides a framework for studying carbohydrate-active enzymes and other viral proteins across the virosphere.

## Methods

### Data collection

The viral gene and genome data were downloaded from the KEGG database on February 14, 2025. KEGG was selected as the primary data source because it provides stable gene identifiers and direct links to KEGG Orthology (KO) assignments and other curated functional information. Broader resources such as NCBI Virus may be incorporated in future expansions, whereas the present version prioritizes stable identifiers and integration with KO annotations. The mapping profiles of KEGG IDs to other accessions, along with the Virus-Host Database^24^ and the viral metadata resource (VMR) MSL39.v4^25^ from International Committee on Taxonomy of Viruses (ICTV), were used. Viral genes are categorized into two parts: viral genes and viral mature peptides. The statistics of viral genes in the KEGG database can be accessed at: https://www.kegg.jp/kegg/genome/virus.html.

### Polyproteins processing

Polyproteins are challenging to identify due to their complex structure and diverse annotations. To generate a set of high-confidence polyprotein labels, we employed a combined approach. First, we scanned gene descriptions and labeled any gene as a polyprotein if its annotation contained the keyword “polyprotein.” Second, we performed BLAST searches of all viral proteins against the KEGG mature peptide database, which includes curated mature peptides cleaved from polyproteins in selected viruses. For each viral gene, we quantified its nucleotide length and its relative length compared to the corresponding genome. Notably, some genomes in the dataset were incomplete, segmented, or represented only coding sequences. We observed that genes (N = 1,048) identified by both the keyword-based and BLAST-based approaches were typically long and accounted for at least 14% of the total genome length. These genes were used as high-confidence polyprotein labels to supervise downstream segmentation analysis and were designated as positive labels. In parallel, genes that were not identified by either method were treated as negative labels for subsequent performance evaluation. For a balanced performance assessment, 2,000 sequences longer than 2,000 amino acids were randomly selected.

All protein-coding genes were then analyzed using hmmscan^32^ against two VOG databases: 1) the public VOGDB^26^ and 2) a beta version of VirGenes developed in this study. HMM hits were mapped to gene sequences to construct per-residue coverage profiles, where each amino acid position was assigned a hit count. To identify potential cleavage regions, we applied Savitzky–Golay filtering to smooth the coverage profile, followed by detection of local minima as candidate cut points.

A gene was marked as a putative polyprotein if its smoothed coverage profile yielded at least one cut point, resulting in segmentation into two or more regions. These predictions were evaluated against the initial labels to optimize the segmentation parameters. The best performance (highest F1 score) was achieved using a minimum protein length threshold of 1500 amino acids and the following smoothing parameters: smooth_window = 111, polyorder = 2, and prominence = 1.5. This configuration was subsequently applied to all proteins to systematically identify and segment putative polyproteins. In total, 4,958 sequences were labeled as polyproteins and segmented into 26,738 regions for the subsequent clustering step. Additionally, 51,494 sequences were excluded from the polyprotein detection step due to the absence of any hmmscan hits. A total of 703,080 sequences were included in the clustering step.

### Viral Sequence Clusters (VSCs)

The first step in constructing VirGenes is to construct gene cluster profiles based on the sequence alignment results. The gene clustering includes two steps: 1) a global pairwise gene alignment and 2) cluster based on the graph approach. To reduce the influence from partial alignment (namely only fragments of sequences are aligned to the other one), we performed a correction on the identity as follows:

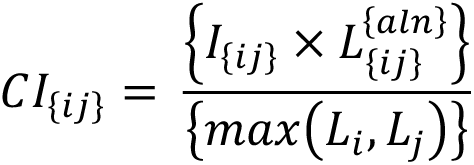

Then the corrected identity was used as the weight of edges between viral genes. The generated network was cut using Markov Clustering Algorithms (MCLs)^33^.

The best combination of sequence alignment tools and inflation index of MCLs were selected according to an assessment test. In brief, all KOs assigned to viral genes were used as the positive labels and the homogeneity between the VSCs in this step and KO labels were calculated. Four indices, Adjusted Rand Index, Normalized Mutual Information, Completeness Score, and Homogeneity Score, were used in the assessment. The combination of MMseqs2^34^ and inflation index of 2 was selected. After this step, 63,216 VSCs were generated, and, among these, 42,297 VSCs contained more than three viral genes.

### Refinement of VSCs

We performed three methods to remove the “bad” members of each VSC. The first approach is based on the sequence alignment that we redo the alignment using DIAMOND^35^ within each VSC, independently. The intra-cluster centrality was calculated for each member gene. Then the outliers for each cluster were removed. Genes with centrality values below the defined cut-off were removed. The cut-off was determined using both the 1) mean - 2 SD criterion and 2) the Q1 - 1.5 × IQR criterion. Similar treatment was done based on the protein length.

Then we performed multiple sequence alignment using MAFFT^36^ and HMM and generated HMM profiles for all VSCs using HMMER. Each gene was assigned with a best fit HMM profile using hmmscan. If a gene was assigned with the cluster it belonged to, it will be retained. If a gene was assigned with a cluster different from the one it originated, the gene will be removed and the HMM profile of this cluster will be reconstructed. This step runs iteratively until all the genes have the best fit model of a cluster it comes from.

### Remote homologous clustering

We first curated VSC groups based on HMM profiles constructed for each VSC. Using the resulting clean 42,297 MSA profiles, we built an HHblits^37^ database and performed an all-against-all HHsearch analysis among VSCs. Significant VSC–VSC relationships were retained using an E-value threshold of 5E-2.

This procedure identified 3,063 connected components in the resulting similarity network, among which the largest component (component_1) contained 15,794 VSCs. To further resolve higher-order structure, we applied an independent Louvain community detection algorithm^38^ to the network. In total, 3,898 viral higher-rank clusters (VRCs) were defined.

### Annotation of clusters

VSCs were annotated by summarizing gene-level functional annotations within each sequence cluster. Candidate descriptions were grouped across labels from each tool, and the representative description was selected using a majority-rule principle based on the fraction of member genes supporting each description. VRC annotations were then summarized from the VSC-level annotations and common domain information. The resulting VSC and VRC annotations were further refined by manual curation to merge synonymous labels.

### Phylogeny of VSCs

Multiple sequence alignments were generated using MAFFT and trimmed with trimAl^39^ using a gap threshold of 0.1. Phylogenetic trees were reconstructed with IQ-TREE v2.2.0^40^ using the -m TEST option, with substitution models automatically selected based on the Bayesian Information Criterion (BIC). For phylogenetic analyses, viral-origin sequences in the CAZyme database were removed and replaced with the CAZyme-VSC sequences identified in this study.

### Protein structures of VSCs and CAZymes

A recent study comprehensively predicted viral protein structures using AlphaFold, and we downloaded the corresponding dataset^13^ for integration into VirGenes. In parallel, we scanned the Protein Data Bank (PDB)^41^ and extracted experimentally determined structures corresponding to VSC members. For VSCs that lacked any representative in either the AlphaFold dataset or the PDB, we selected a representative protein defined as the member with the highest intra-cluster centrality based on intra-group sequence alignment. The structures of these representative proteins were then predicted using AlphaFold^42^.

In total, 97,270 protein structures were obtained and incorporated into VirGenes. An all-against-all structural comparison was then performed using Foldseek^43^. Significant structural similarities were retained using an E-value threshold of 5E-2. For each significant structure pair, the minimum TM-score among query TM-score (qTM-score), target TM-score (tTM-score), and alignment TM-score (alnTM-score) was selected as the edge weight for network construction. Using this structural similarity network, connected components were first identified following the same definition applied in the VRC analysis. Subsequently, Louvain community detection was performed to further resolve higher-order structural relationships.

### Detection of CAZyme-encoding viral genes

Detection of CAZyme-encoding viral genes was primarily based on BLAST^44^ searches against the CAZy database^1^, retaining hits with an E-value ≤ 1E−5, at least 60% sequence similarity, and at least 60% coverage. These searches were complemented by hmmscan against the dbCAN HMM database^17^, applying a stringent cutoff of E-value ≤ 1E−15 and HMM coverage ≥ 0.35. Because CAZy/dbCAN primarily captures CAZyme families, this search does not comprehensively detect enzymes involved in nucleotide-sugar biosynthesis or nucleotide-sugar modification.

To identify VSCs with CAZyme-like folds, representative structures of VSCs were compared against representative structures of known CAZyme families using Foldseek. Significant VSC–CAZyme structural matches were retained using an E-value threshold of 5 × 10−2. The minimum of the qTM-score, tTM-score and alnTM-score was used as the structural similarity score.

### Binding-pocket prediction and molecular docking

Binding pockets were predicted from the AlphaFold3 models^45^ using P2Rank v2.5.1^46^ through PrankWeb^47^, and the highest-ranking pocket located in the central groove of each β-propeller was selected for docking. Molecular docking was performed using AutoDock Vina v1.2.7^48^ with a 20 × 20 × 20 Å search box centered on the predicted pocket. β-1,3-Mannobiose was used as the positive-control ligand^49^, whereas β-1,4-mannobiose, cellobiose, and maltose were used as control disaccharides. Each docking experiment was performed using three random seeds, and the predicted binding energies are reported as the mean ± SD of the best-scoring poses. Docking poses were visualized using PrankWeb.

### VirGenes updating

To incorporate newly available viral protein sequences while maintaining the stability of the VirGenes hierarchical framework, we implemented a streamlined update procedure briefly as follows. First, newly added viral proteins were identified by comparing the latest KEGG viral gene dataset with the previously released VirGenes data. These new sequences were extracted and screened against the HMM profiles representing all existing VSCs. Matches passing predefined score and coverage thresholds were assigned to their corresponding VSCs.

Proteins without significant hits were subjected to de novo clustering with previous singleton genes using MMseqs2 and MCL to form new VSCs as the initial frame, ensuring that previously unrepresented gene families were integrated into the database. Finally, newly assigned and newly formed VSCs were merged with the existing VirGenes hierarchical structure (VSC–VRC–VFC), and functional annotations were propagated accordingly. This procedure enables continuous expansion of VirGenes while preserving cluster compatibility with earlier versions.

This automated workflow supports routine incremental updates of VirGenes to keep the accession IDs of VSCs identical. However, major updates that involve redefining cluster boundaries or restructuring higher-level ortholog groups will be performed manually to ensure biological consistency.

## Supplementary material

The supplementary material includes five supplementary figures and one supplementary table. Figure S1 provides an overview of the VirGenes hierarchical workflow. Figure S2 describes the identification and segmentation of viral polyproteins. Figure S3 shows the distribution of sequence identity within VSCs. Figure S4 evaluates the consistency of CAZyme annotations within VSCs. Figure S5 illustrates the organization of CAZyme-associated VSCs at the VRC level. Table S1 summarizes the parameters used for viral gene clustering.

## Data Availability Statement

VirGenes and the viral gene clusters generated in this study are freely accessible at https://www.genome.jp/vogdb/. The viral gene and genome data analyzed in this study were obtained from KEGG, and the associated viral taxonomy and host information were obtained from the ICTV Virus Metadata Resource and Virus–Host DB, respectively. CAZyme annotations were based on CAZy and dbCAN. Additional data supporting the findings of this study are available from the corresponding author upon reasonable request.

## Supporting information

Spplementary Figures and Table

## Acknowledgments

This work was supported by a research grant from HFSP (Ref.-No: RGP011/2024; https://doi.org/10.52044/HFSP.RGP0112024.pc.gr.194158), the NBDC Database Integration Coordination Program JPMJND2203 of the Japan Science and Technology Agency, the Collaborative Research Program of the Institute for Chemical Research, Kyoto University (2026-27), and JSPS KAKENHI (22H00384). Computational analyses were performed using the Supercomputer System of the Institute for Chemical Research, Kyoto University.

## AI Disclosure Statement

Generative artificial intelligence tools were used to assist with language editing and selected coding tasks. All generated text and code were reviewed and verified by the authors. The authors take full responsibility for the scientific content, analyses, interpretations, and conclusions presented in this manuscript.

