## Supplementary material for "A hierarchical orthology framework reveals viral carbohydrate-active genes across the global virosphere": Spplementary Figures and Table

### Running Title: VirGenes reveals viral carbohydrate-active genes

Lingjie Meng[1], Ruixuan Zhang[1], Cristina De Castro[2], Ikuo Uchiyama[3], Minoru Kanehisa[1], Hiroyuki Ogata[1,*]

1. Institute for Chemical Research, Kyoto University, Gokasho, Uji, Japan
2. Department of Chemical Sciences, University of Naples Federico II, Naples 80126, Italy
3. National Institute for Basic Biology, National Institutes of Natural Sciences, Okazaki, Japan

[*] *Corresponding author*

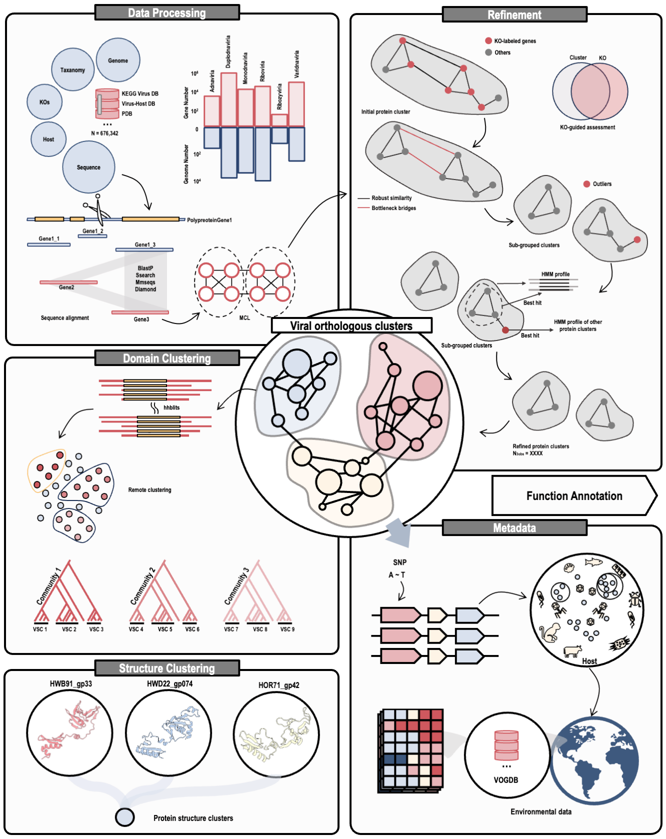


**Supplementary Figure 1. Overview of the VirGenes hierarchical workflow.**
Schematic summary of viral gene collection, polyprotein processing, sequence clustering, profile-based remote clustering, structure-based fold clustering and functional annotation.

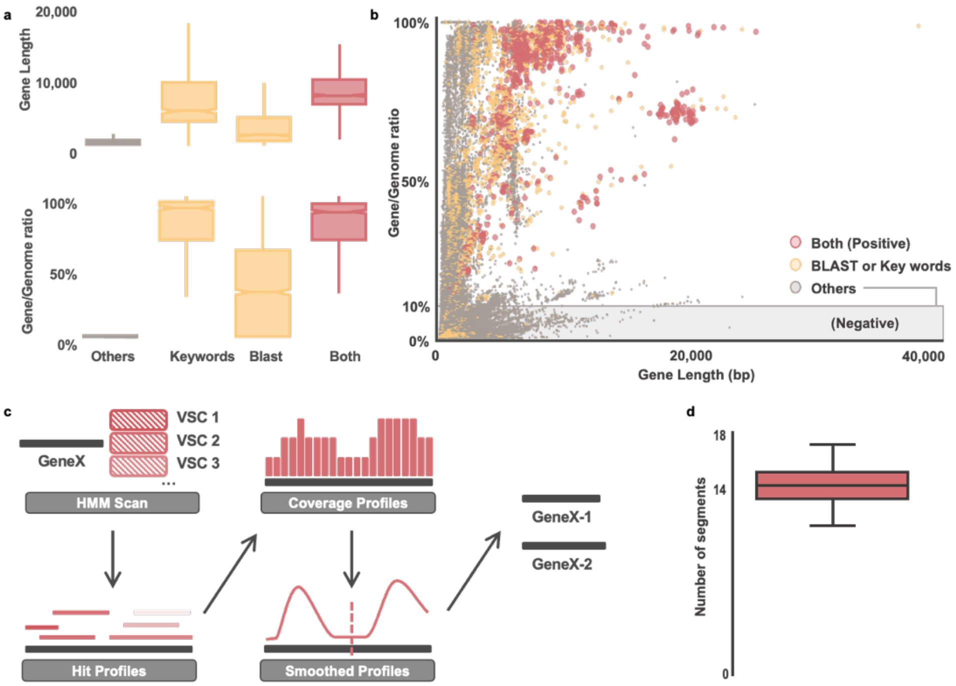


**Supplementary Figure 2. Identification and segmentation of viral polyproteins.**
a, Gene length and gene/genome ratio across polyprotein-evidence groups. b, Scatterplot of gene length versus gene/genome ratio. c, Profile-based workflow for detecting putative cleavage sites in polyproteins. d, Distribution of predicted segment numbers per polyprotein.

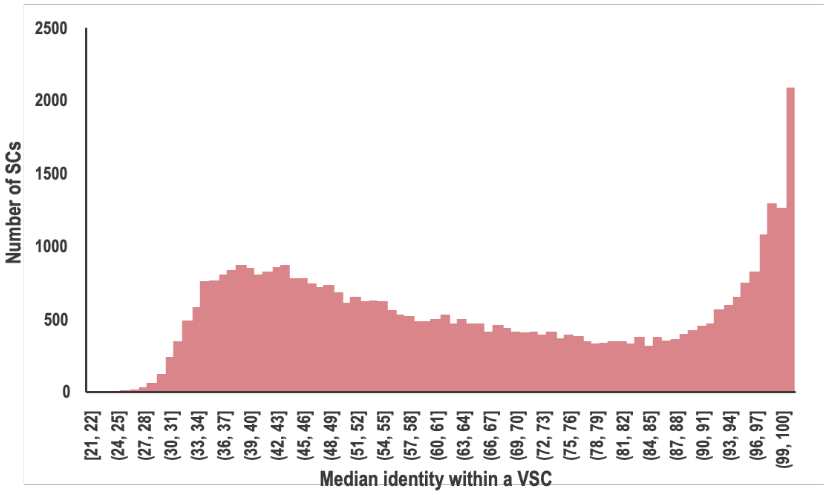


**Supplementary Figure 3. Distribution of median identity of VSCs.** Histogram showing the median pairwise sequence identity within each viral sequence cluster (VSC).

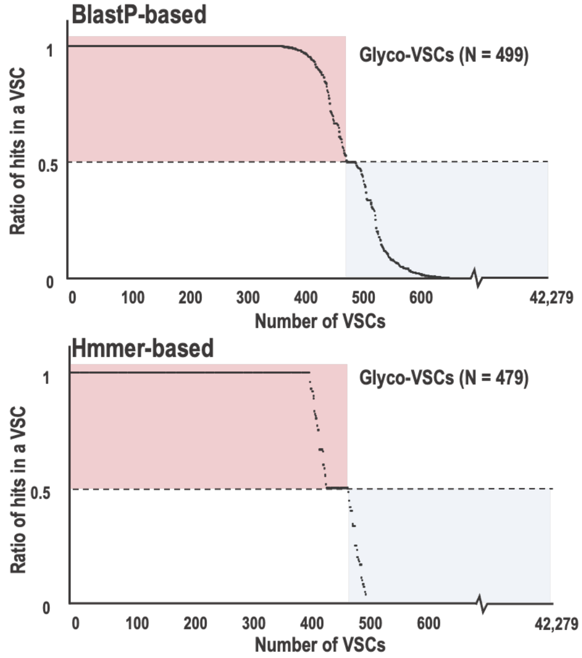


**Supplementary Figure 4. Detection of CAZyme-encoding genes in VSCs.**
BLASTP- and HMMER-based evaluation of CAZyme annotation consistency within VSCs.

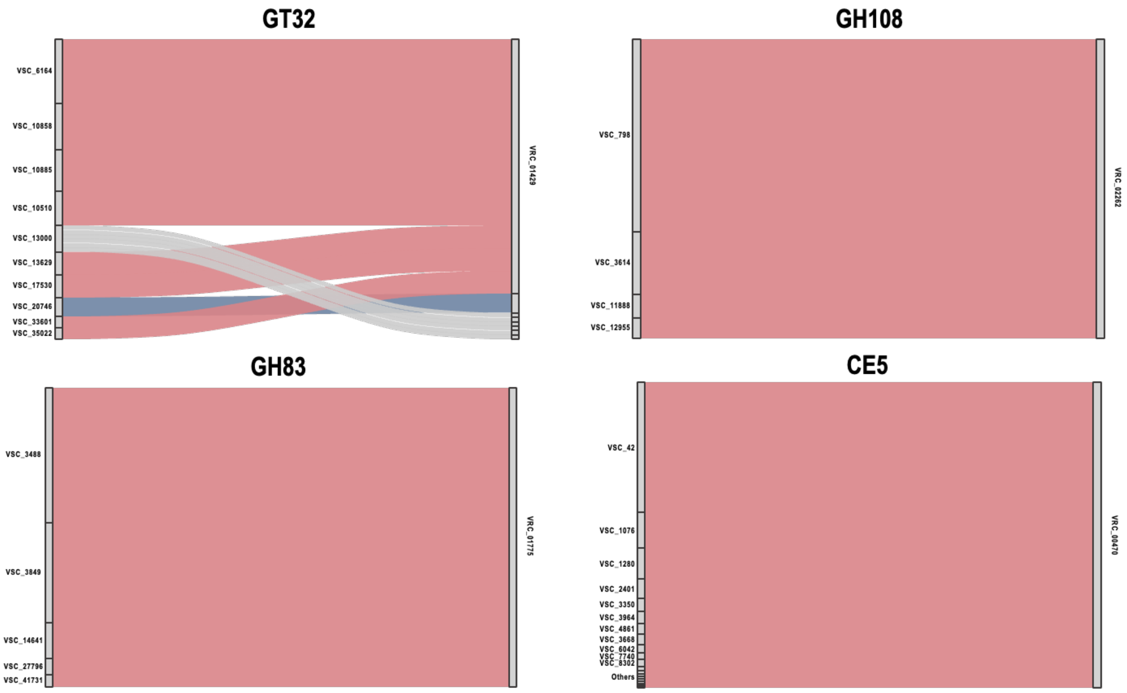


**Supplementary Figure 5. Detection of CAZyme-associated genes in VRCs.**
Examples showing how CAZyme-associated VSCs are grouped into higher-order viral remote clusters (VRCs) by profile-level similarity.

**Supplementary Table 1. Evaluation of sequence clustering parameters.**

| Method | Inflation | ARI | NMI | Homogeneity | Completeness | V-measure |
|---|---|---|---|---|---|---|
| BlastP | i2 | 0.7665 | 0.9596 | 0.961 | 0.9583 | 0.9596 |
| | i3 | 0.774 | 0.9594 | 0.9701 | 0.949 | 0.9594 |
| | i4 | 0.7724 | 0.9588 | 0.9745 | 0.9435 | 0.9588 |
| | i5 | 0.764 | 0.9572 | 0.9769 | 0.9382 | 0.9572 |
| Diamond | i2 | 0.4013 | 0.9177 | 0.9954 | 0.8513 | 0.9177 |
| | i3 | 0.3347 | 0.9043 | 0.9974 | 0.8271 | 0.9043 |
| | i4 | 0.2947 | 0.8941 | 0.9981 | 0.8097 | 0.8941 |
| | i5 | 0.2673 | 0.8859 | 0.9984 | 0.7962 | 0.8859 |
| MMseqs2 | i2 | 0.7829 | 0.9617 | 0.9853 | 0.9393 | 0.9617 |
| | i3 | 0.769 | 0.9582 | 0.9882 | 0.93 | 0.9582 |
| | i4 | 0.7609 | 0.9563 | 0.9898 | 0.9249 | 0.9563 |
| | i5 | 0.7579 | 0.9551 | 0.9921 | 0.9208 | 0.9551 |
| Ssearch | i2 | 0.7413 | 0.955 | 0.9518 | 0.9582 | 0.955 |
| | i3 | 0.7575 | 0.956 | 0.9629 | 0.9492 | 0.956 |
| | i4 | 0.7582 | 0.9555 | 0.9687 | 0.9427 | 0.9555 |
| | i5 | 0.7509 | 0.9533 | 0.972 | 0.9353 | 0.9533 |
